# Deciphering Cargo Contents in Extracellular Vesicles of *Candida haemulonii* var. *vulnera*

**DOI:** 10.1101/2024.08.12.607614

**Authors:** Bianca T. M. Oliveira, Tamires A. Bitencourt, Patrick W. Santos, Antônio D. Pagano, André M. Pessoni, Caroline P. Rezende, Renan A. Piraine, Ana P. Masson, Vitor Faça, Vinicius F. Campos, Lysangela R. Alves, Arnaldo L. Colombo, Fausto Almeida

## Abstract

*Candida haemulonii* comprises a group of pathogenic fungi known for their resistance to primary antifungal treatments. Infections caused by these pathogens present substantial challenges due to the difficulties in accurate identification. Extracellular vesicles (EVs) released by these fungi play a critical role in the pathogen-host interaction, potentially influencing antifungal resistance and virulence. Previous research by our group indicates that EVs contain immunogenic particles capable of impacting the host’s immune response. Understanding the composition of these EVs is crucial for elucidating the mechanisms underlying resistance and virulence in *C. haemulonii* var. *vulnera*. This study aims to investigate the contents of EVs from *C. haemulonii* var. *vulnera* using proteomic and microRNA sequencing tools, providing insights into their role in adaptation, survival, and the progression of infections. Our findings reveal key proteins transported by EVs, including BMH1, TEF1, CDC19, and PDC11. These proteins are involved in various cellular processes, such as the alteration of cell wall structure, biofilm formation, and facilitation of morphological changes, among others. Additionally, we observed that miRNA-like molecules transported within EVs are linked to the electron transport chain and regulation of the citric acid cycle, which are metabolic processes associated with virulence factors and rapid adaptation to diverse hosts or environments. In this context, our findings provide a novel perspective on fungal EVs, highlighting their potential as targets for therapies. Therefore, these vesicles may reflect the expression levels of regulatory molecules crucial for the survival, pathogenicity, and virulence of *C. haemulonii* var. *vulnera*.

**IMPORTANCE:** The study of *Candida haemulonii* complex holds substantial clinical significance due to its notable resistance to conventional antifungal therapies and the complex challenges inherent in its specific identification. This research focuses on cargo of EVs released by these fungi, which play an essential role in pathogen-host interactions, influencing fungal pathogenicity. EVs contain immunogenic particles that can modulate the host’s immune response. Proteomic and microRNA analyses of EVs from *Candida haemulonii* var. *vulnera* have identified key proteins and miRNAs involved in cellular processes such as metabolic adjustment, biofilm formation, and modulation of cytoplasmic functions. These components are essential for the adaptation, survival, and progression of infections. This study offers novel insights into fungal EVs, underscoring their potential as targets for therapeutic intervention. By elucidating the mechanisms underlying the rapid adaptation of *Candida haemulonii*, the research enhances our understanding of the pathogenicity of this emerging yeast.

## INTRODUCTION

*Candida haemulonii* represents a complex group of closely related emerging fungal strains, including *Candida haemulonii sensu stricto*, *C. haemulonii* var. *vulnera*, *C. duobushaemulonii*, *C. vulturna* [1–3]. Accurately identifying these pathogens is challenging due to their phenotypic resemblance to other *Candida* species [4, 5]. These species are known to cause infections in immunocompromised individuals, such as bloodstream, urinary tract, and wound infections[6, 7]. Global reports from healthcare settings have underscored their resistance to multiple antifungal agents, contributing to elevated mortality rates [8, 9].

During infection, effective control of fungal growth and mitigation of yeast-induced damage are crucial for delaying disease progression in the host [10, 11]. In this context, extracellular vesicles (EVs) play a pivotal role in the interaction between the pathogen and host [12, 13]. EVs are bioactive molecules enclosed in a lipid bilayer that facilitate intercellular communication and exchange of genetic material upon release from fungal cells, thereby influencing rapid fungal adaptation and survival [14–16]. Consequently, EVs may contribute to antifungal resistance by transferring virulence factors, thereby influencing environmental regulation and pathological processes [17]. Moreover, these EVs serve as essential messengers in bidirectional communication, transmitting biological information and participating in the intricate fungus-host interaction[18].

Our research group previously demonstrated that EVs from *C. haemulonii var. vulnera* are recognized by murine macrophages, triggering an oxidative burst [19]. This finding indicates that the content of these EVs contains immunogenic particles that are crucial during the evolution of infection [20–22]. EVs derived from *C. haemulonii* have the potential to enhance the species’ pathogenicity due to their abundant proteins and immunogenic nucleic acids, characteristics not previously described for this strain.

Understanding the content of C. *haemulonii EVs* is essential for comprehending the strain’s multidrug resistance and pathogenicity [23–26]. Integrative proteomic and transcriptomic studies of these fungal EVs can elucidate the intrinsic molecular mechanisms involved in host-pathogen interactions, multidrug resistance, and fungal virulence [27–29].

Proteomics has been consistently utilized to uncover the diverse array of proteins by EVs and their pivotal roles in critical inter- and intracellular communication events throughout infection progression [30–32]. Moreover, identifying immunogenic proteins transported by EVs in *Candida* species enhances our understanding of these molecules’ involvement in stress responses and antifungal mechanisms [33–35]. For instance, it has been observed that EVs from *C. auris* are integral to modulating host cellular defense mechanisms, impacting adhesion to epithelial cells, macrophage-mediated cytotoxicity, and dendritic cell activation[25]. In contrast, EVs from *C. albicans* predominantly contain cell wall-related proteins, secreted enzymes crucial for pathogenesis and nutrient acquisition, as well as cytoplasmic proteins[36, 37].

On the other hand, RNA-Seq investigates the composition of RNAs in a specific biological state, offering a comprehensive view of genetic information exchange between cells [38, 39]. Notably, during infection, the abundance of miRNA-likes transported by EVs serves as a survival strategy [40, 41]. These miRNA-likes function as modulators of gene expression, potentially activating or repressing the translation of target messenger RNA molecules [42, 43]. Previous studies, such as those involving the gastrointestinal parasite *Heligmosomoides polygyrus,* have documented how miRNA-likes exported by EVs neutralize host responses [44]. In another context, plant EVs have been shown to accumulate at cellular levels in phytopathogenic fungi [45]. Through small RNA (sRNA) profiling, researchers identified the downregulation of target genes associated with reduced fungal virulence [46].

In this context, investigating the contents of *Candida haemulonii* EVs is crucial for understanding the biomolecules carried by the EVs of this emerging species, known for its virulence and multidrug resistance, which poses significant challenges in hospitals and intensive care units. In the present study, we present, for the first time, a comprehensive characterization of EVs composition from *Candida haemulonii* var. *vulnera* using omics tools such as proteomics and RNA-seq. This approach allows us to unveil the molecular makeup of EVs from this emerging yeast and offers insights into the development of effective antifungal strategies against this potentially dangerous species.

## MATERIALS AND METHODS

### Fungal Strains and Growth Conditions

The *Candida haemulonii* var. *vulnera* ATCC 1112/2016 strain was cultivated at 30°C in 4% Sabouraud Dextrose Agar (SDA) medium adjusted to pH 5.6 for 48h. Subsequently, four fresh colonies were transferred into 5ml of Sabouraud broth and cultured at 30°C with shaking at 150 rpm. Following this, an experiment was conducted to isolate EVs [19].

### Production and Purification of EVs and Nanoparticle Tracking Analysis (NTA)

The isolation of EVs from *C. haemulonii* var. *vulnera* followed a methodology adapted from Vallejo *et al*. [47] for *Paracoccidioides brasiliensis.* Initially, cells and debris were removed by sequential centrifugation steps at 5000*×g* and 15,000*×g* for 15min to isolate EVs. Subsequently, the supernatants were concentrated using an Amicon ultrafiltration system with a 100kDa cutoff (Millipore, Billerica, MA, USA) and filtered through a 0.45µm membrane (Sigma-Aldrich, St. Louis, MO, USA). The resulting concentrated supernatant was then subjected to ultracentrifugation at 100,000*×*g at 4°C for 1h. The pellets obtained after ultracentrifugation were collected and resuspended in ultra-pure water (Sigma-Aldrich) and supplemented with a 10x protease inhibitor cocktail (PIC) (Sigma-Aldrich) at 0.2 Proteomic analysis of EVs from *C. haemulonii* Protein extraction from yeast cells and EVs

Protein extraction followed an adapted protocol for *C. albicans* [37]. Approximately 40 mg of *C. haemulonii* var. *vulnera* pellets were resuspended in ultrapure water in conical tubes for yeast protein extraction. The suspensions underwent two sequential centrifugations at 3000*×g* for 5min each, with the supernatant discarded after each centrifugation. The resulting pellet was then resuspended in 300µl of lysis buffer (50mM TrisCl pH 8, 150mM NaCl, 1% Triton x100, 1xPIC) and incubated under agitation at 150 rpm at 30°C for 10 min. Subsequently, the samples underwent sonication in 5 cycles of 30 seconds each, with 1min on ice between cycles, using an amplitude of 60%. The sonicated samples were centrifuged at 7000*×g* for 5 min at 4°C, and the resulting supernatant was transferred to a new microtube.

Following homogenization, an aliquot of 200µl from yeast protein was quantified using 96-well plates, while the remaining sample was stored at -80°C. For protein extracting from EVs, the total amount of EVs was concentrated by ultracentrifugation at 100,000*×g* for 1h at 4°C. The supernatant was discarded, and the EV pellet was resuspended in Tris-SDS buffer (20 mM Tris.HCl pH 8) supplemented with 0.2% PIC (Sigma-Aldrich). The solution was heated at 95°C for 5min using a dry bath.

Subsequently, microdilution tubes were sonicated for 1min at 60% amplitude with a pulse of 10 seconds on and 10 seconds off. Finally, the microtubes were centrifuged at 15,000 rpm for 20 min at 4°C, and the resulting supernatant containing EV proteins was transferred to new microtubes and stored at -80°C.

### Measurement and digestion of yeast and EV proteins

The concentration of *C. haemulonii* var. *vulnera* yeast proteins were quantified using the Lowry method, following the manufacturer’s guidelines (Bio-Rad Laboratories, Hercules, CA, USA). Measurements were conducted on an adjustable microplate reader (Bio-Rad iMark Microplate Reader). For EV proteins, quantification was performed using the bicinchoninic acid (BCA) method (Sigma-Aldrich, USA). The methodology outlined by Ferreira *et al*. [48] was followed for further analysis. Proteins were separated by SDS-PAGE using 4–20% Mini-PROTEAN TGX Precast Protein Gels (Bio-Rad Cat # 4561093). Each gel lane was divided into 4 equal-sized pieces, and *gel* digestion was performed individually for each piece. SDS and dye were removed using NH_4_HCO_3_ (50mM) with 50% acetonitrile (ACN), followed by a wash with pure ACN. The gel slices were dried in SpeedVac (Savant), rehydrated with NH_4_HCO_3_ (100mM) containing 0.6µg of trypsin (Promega) for approximately 30min followed by adjusting the volume in the tube with NH_4_HCO_3_ (100mM) to fully cover band pieces. The digestion was carried out at 37°C for 16–18h. After digestion, peptides were extracted from the gel using consecutive washes with 0.1% formic acid solutions containing ACN at 50%, 70%, and 100%. The extracted peptides were transferred to clean low-protein binding microtubes and dried using SpeedVac (Thermo Scientific). The dried samples were resolubilized in 5% ACN containing 0.1% formic acid and desalted using C_18_ Zip Tips™ columns (Sulpelco, Sigma) following the manufacturer’s instructions. Triplicate samples of yeast proteins, *C. haemulonii* var. *vulnera* EVs proteins, and a BSA control were submitted to the Analytical Instrumentation Center of the University of São Paulo (Analytical Center - Institute of Chemistry) for Nano LC-MS Mass Spectrometry (Complex samples - gradient 240min) using the MaXis - QTOF - High-resolution equipment.

### Proteomic data analysis

Following the methodology outlined by Amorim *et al.* [49], with some modifications., raw data were processed using Peaks 8.5 software (Bioinformatics Solutions, Waterloo, Canada) against a database containing the keyword “*Candida*” downloaded from the Uniprot database downloaded in May 2022. Carbamidomethylation was specified as a fixed cysteine modification, while oxidation (M) and amidation were considered variable modifications for methionine and glutamine, respectively. Up to 3 missed cleavages were allowed for tryptic peptides. Mass error tolerances were set to 5 ppm for parent ions and 0.015 Da for fragment ions. A false discovery rate (FDR) of 1

### High-throughput sequencing of miRNA-like

Genetic material was extracted from the fungal mass of *C. haemulonii* var. *vulnera* and its EVs to identify and characterize the RNA content. Total RNA was extracted from three biological replicates of yeast and three biological replicates of EVs cultured on SDA for 48h at 30°C. Cells were harvested by centrifugation, immediately frozen in liquid nitrogen, and lysed using a lytic enzyme to isolate total RNA. EVs were purified as described in section 2.2 (with 20 extractions per sample). RNA extraction was performed using Trizol (Invitrogen), followed by treatment with RNase-free DNase I (Fermentas), and purification using the miRNeasy Mini Kit (Qiagen Biotecnologia Brasil Ltda, São Paulo, SP, Brazil), following the manufacturer’s protocol, to yield a final concentration of 1 µg RNA per sample.

The extracted miRNA was quantified using the Qubit 3.0 Fluorometer (Life Technologies), and its integrity was verified using the Agilent 2100 Bioanalyzer system. To assess sample integrity, an induced degradation test was conducted with RNAse to ensure quality. Libraries were prepared and sequenced at the Cancer Center - Institute for Cancer Research - IPEC (Paraná, Brazil) using the QIAseq miRNA Library Kit Handbook and QIAseq miRNA 96 INDEX IL preparation kit – both from QIAGEN, following the manufacturer’s protocols. Subsequent clustering and sequencing were performed on the Illumina NextSeq 500 system. Paired-end reads were multiplexed and analyzed against reference genes of *C. haemulonii* var. *vulnera* using Kallisto software (version 0.43.1).

The identification and functional enrichment of miRNA-like sequences in *C. haemulonii* var. *vulnera* were conducted using QIAGEN CLC Genomics Workbench (version 24.0.1). Bioinformatics analyses of miRNA-seq data were performed separately for yeast and EV libraries. Initially, raw sequencing reads underwent preprocessing to remove adapter sequences (QIAseq miRNA NGS Adapters: 5’ GTTCAGAGTTCTACAGTCCGACGATC; 3’ AACTGTAGGCACCATCAAT), oversized insertions, ambiguous nucleotides, and low-quality sequences. The miRNA transcriptome was then assembled and mapped using *Candida albicans* as a reference organism (GenBank Genome Assembly ASM18296v3, Accession No. GCA_000182965.3). Mapping parameters included a match score of 1, mismatch cost of 2, linear gap costs for insertions and deletions, and specific criteria for length and similarity: insertion cost of 3, deletion cost of 3, length fraction of 0.5, similarity fraction of 0.8, and minimum seed length of 15. To characterize miRNA-like sequences, mapped reads were annotated based on miRNA seed sequences from the miRBase database (version 22.0), encompassing 10.903 miRNA-like from 271 organisms. Expression levels of miRNA-like sequences in *C. haemulonii* var. *vulnera* were visualized in a heatmap using Euclidean distance for distance measure, complete linkage for linkage criteria, and a feature count threshold. For functional enrichment analysis, sequencing reads were mapped against a fungal reference genome of *Saccharomyces cerevisiae* strain S288C (GenBank Genome Assembly R64, Accession No. GCA_000146045.2). Variant tracks were extracted to retrieve Gene Ontology (GO) annotations, encompassing molecular functions, biological processes, and cellular compartments. The complete list of enriched GO functional pathways can be found in the supplementary materials (Supplementary).

### Interactomics Analysis: Network-Based Visualization of Molecular Interactions Between miRNA-like and Enriched Genes

The QIAGEN CLC Genomics Workbench (version 24.0.1) was employed to identify target genes associated with enriched GO terms influenced by *C. haemulonii* miRNA-like, with a significance threshold set at *p-*value < 0.05. Interactomics analysis was performed separately for both yeast and EVs libraries. Biological networks of genes enriched by *C. haemulonii* miRNA-like were constructed using the STRING (Search Tool for the Retrieval of Interacting Genes/Proteins) database (https://string-db.org/), with co-expression selected as the interaction source. The reference organism used was *Candida albicans* (var. SC5314, NCBI taxonomy Id: 237561). Interactomes from STRING were retrieved and visualized in Cytoscape (version 3.10.2)[49]. To identify clusters (highly interconnected regions) with the gene-gene networks, the Cytoscape app Molecular Complex Detection (MCODE)[50] was utilized. Additionally, the Cytoscape plugin cytoHubba[51] was employed to rank the hub genes based on Degree, a topological analysis method. In the visualization, genes regulating the enriched GO terms influenced by *C. haemulonii* miRNA-like are represented as green nodes, while hub genes are highlighted as red nodes. The thickness of the edges indicates the strength of interactions, with a high confidence threshold (0.7) used for the interaction score.

### Statical Analysis

The experiments were carried out in triplicate, with each set of experiments independently performed three times. Statistical analysis was conducted using GraphPad Prism version 8.0.1. Results were expressed as the mean ± standard error of the mean (SEM). To identify differences between the studied values, a one-way analysis of variance (ANOVA) followed by the Dunnet post-test was employed. Statistical significance was considered at *p < 0.05.

## RESULTS

### Proteomic profiling of *Candida haemulonii* var. *vulnera* and its extracellular vesicles

Protein profiles from yeast cells and EVs of *Candida haemulonii* var. *vulnera* 1112/2016 were identified using mass spectrometry and categorized based on their functions and cellular localization. After data processing, we identified 178 proteins in yeast cells and 124 proteins in EVs of *C. haemulonii* var. *vulnera* (Figure 1).

**FIG 1.**
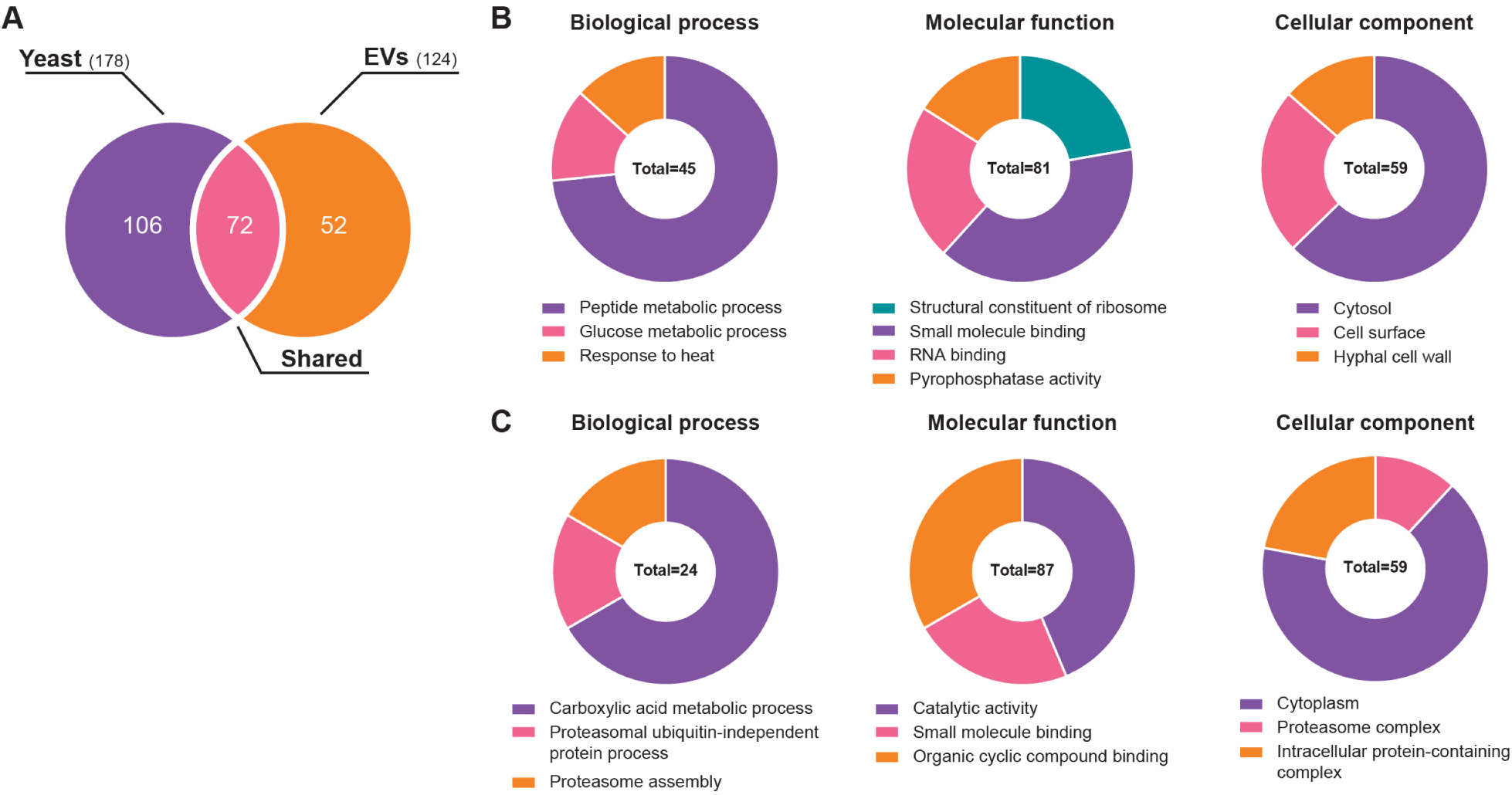
Proteomic profile of proteins found in *Candida haemulonii* var. vulnera 1112/2016 yeast and exclusive proteins transported by extracellular vesicles (EVs) from *Candida haemulonii* var. *vulnera* 1112/2016. A) Venn diagram illustrating the number of proteins identified as either common or exclusive to EVs from *Candida haemulonii* yeast. B) Percentage of proteins common to *Candida haemulonii* cells involved in biological processes, molecular functions, and cellular processes. C) Percentage of proteins exclusive to EVs from *Candida haemulonii* involved in biological processes, molecular functions, and cellular processes Additionally, Gene Ontology was utilized for functional categorization to explore the biological role of EV proteins. Notably, EVs were found to contain active proteasome complexes, which are distinct from the proteins found in yeast cells (Figure 1). This suggests a selective mechanism for protein export via EVs in *C. haemulonii* var. *vulnera.* Our findings highlight an interesting protein profile involved in cellular biological processes and molecular functions (see Supplementary Tables 1 and 2).

### Network of significant protein interactions from the proteomic analysis

Network analysis using the STRING database revealed proteins common to both *C. haemulonii* var. *vulnera* yeast cell and EVs fractions, identifying clusters related to carbohydrate metabolism, cell wall dynamics, yeast-hypha transition in response to heat, and biofilm matrix. This suggests robust interactions among proteins crucial for fungal survival, adaptation, and resistance mechanisms. Furthermore, a comparison of other shared proteins between these fractions highlighted a cluster with significant interactions involving ribosomal proteins and glucose metabolism. This observation underscores the importance of these clusters in metabolic homeostasis, supported by enrichment analysis of biological process GO terms such as “Ribosomal Subunit” and “Glycolytic Process,” among others (see Fig. 2A and 2B).

**FIG 2.**
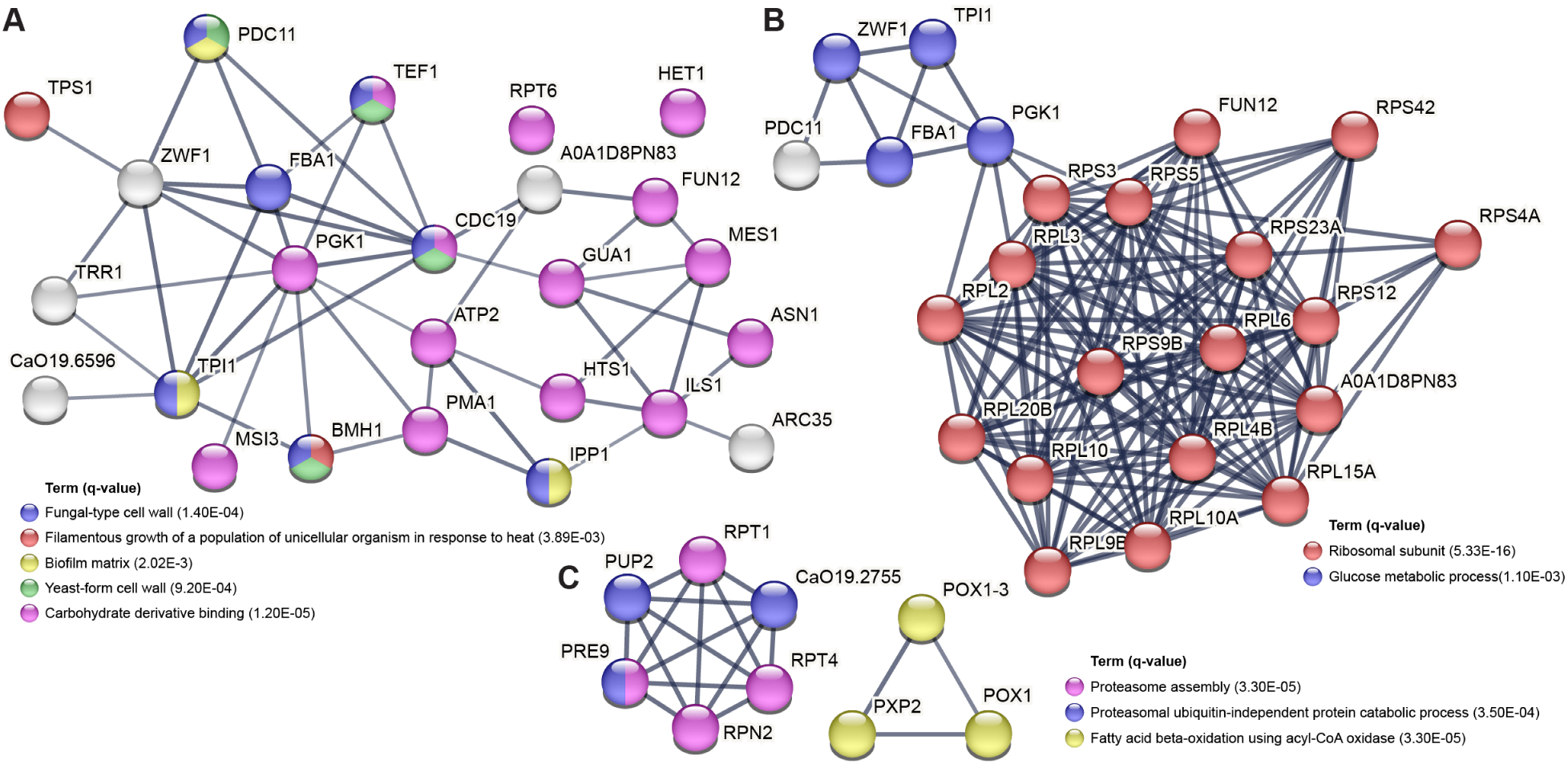
**Interaction networks of proteins from yeast and extracellular vesicles (EVs) from *Candida haemulonii* var. *vulnera***. **A)** Protein-protein interaction network derived from commonly expressed proteins with increased expression in both yeast cells and EVs of *Candida haemulonii* var. *vulnera*. **(B)** Network of protein-protein interactions involving ribosomal proteins with common expression in both yeast cells and EVs of *Candida haemulonii* var. *vulnera*. **C)** Protein-protein interaction network based on proteins showing increased expression, specifically in EVs of *Candida haemulonii* var. *vulnera*. All protein interactions depicted are statistically significant (p<0.05). Highlighted terms represent relevant biological functions. For the comparison, the STRING database (v.12.0) protein-protein association networks and functional enrichment analyses were used [REF PMID: 36370105]

On the contrary, proteins exclusive to EVs clustered predominantly into two categories. One cluster exhibited significant interactions among proteins involved in the assembly of the proteasome complex, while the other cluster was associated with fatty acid metabolism. Notably, no additional enriched terms were identified (Fig. 2C).

Based on these findings, we investigated four key proteins found in both fractions - EVs and fungal yeast cells. They include: 14-3-3 protein homolog (BMH1), belonging to the 14-3-3 family; Elongation factor 1-alpha (TEF1), a member of the TRAFAC class translation factor GTPase superfamily; Pyruvate kinase (CDC19); and Pyruvate decarboxylase (PDC11), classified under the TPP enzyme family. GO analysis revealed that these proteins are integral to a diverse array of cellular processes. These include metabolic adaptation, cell wall modification, biofilm development, cytoplasmic activities, maintenance of redox equilibrium, ionic interactions, and morphological transitions (see Fig. 3).

**FIG 3.**
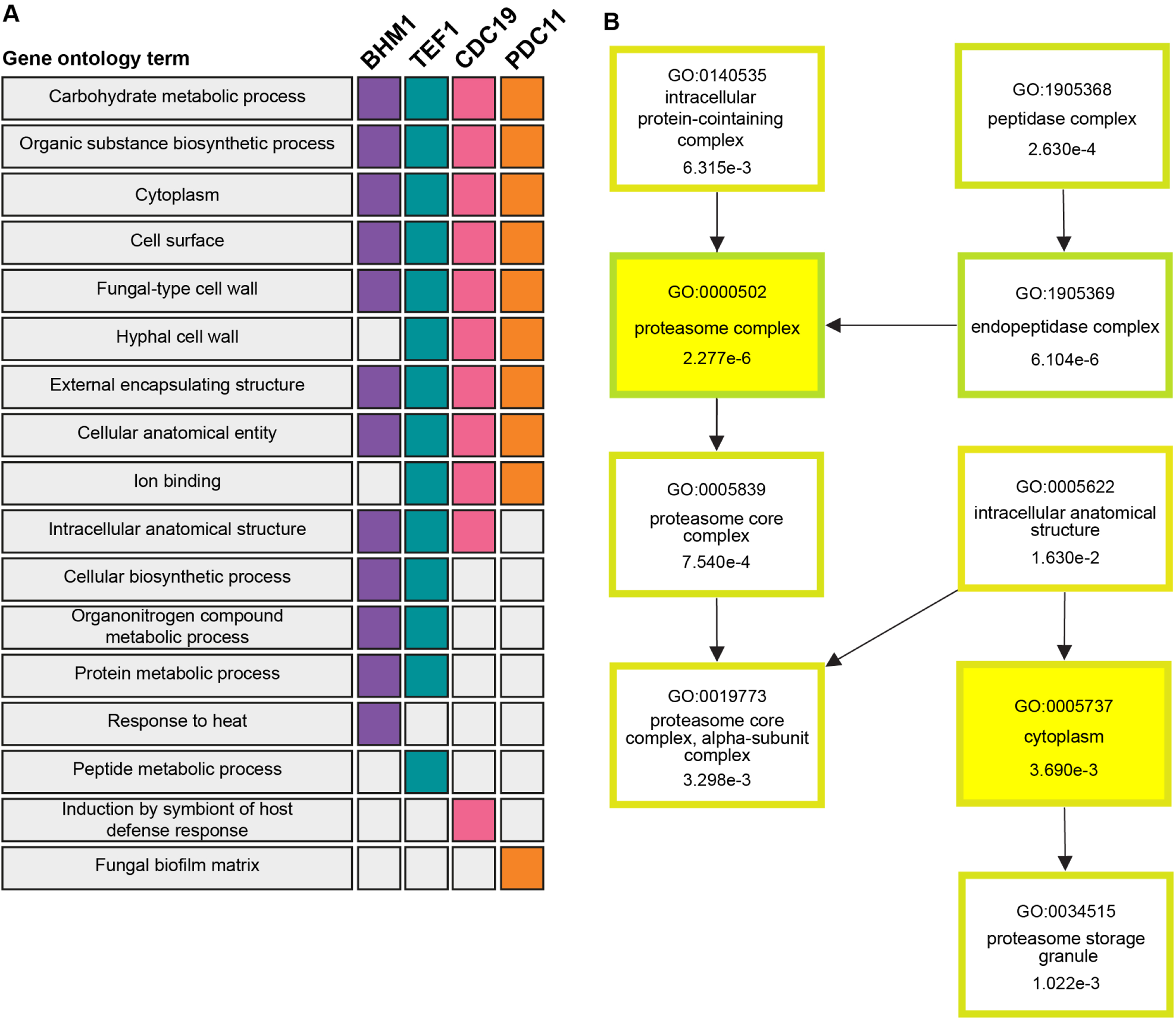
Functions of key proteins in diverse biological processes and transported by the extracellular vesicles (EVs) of *Candida haemulonii* var. *vulnera*. A) Identified proteins: 14-3-3 protein (BMH1); Pyruvate kinase (CDC19); Elongation factor 1-alpha 1 (TEF 1). **B)** Proteins exhibited increased expression in the EVs of *Candida haemulonii* var. *vulnera* significantly contribute to a molecular pathway. The numbers indicate p-values adjusted for list size, where at least 95% of matches are statistically significant.

### Small RNA characterization: miRNA-like sequences for *C. haemulonii* var *vulnera*

MiRNA was extracted from fungal EVs across four independent biological replicates.

Subsequently, RNA samples were enriched in small RNAs (<200 bp) (Supplementary material, Fig. S1).

High-throughput miRNA sequencing of fungal yeast yielded between 3.640.095 and 5.682.755 raw sequencing reads, while fungal EVs produced between 7.483.919 to 28.029.751 raw sequencing reads. Notably, there is no existing literature on miRNA-like specific to fungal species. To identify conserved miRNA sequences among those identified in *Candida haemulonii* var. *vulnera* yeast and EVs, we used a database containing mature miRNA sequences from all characterized organisms to date. We identified 9.843 and 10.406 conserved miRNA-like sequences for the yeast and EVs libraries of *C. haemulonii,* respectively. These results suggest a selective export of miRNA-like by *C. haemulonii* EVs. (Fig. 4).

**FIG 4.**
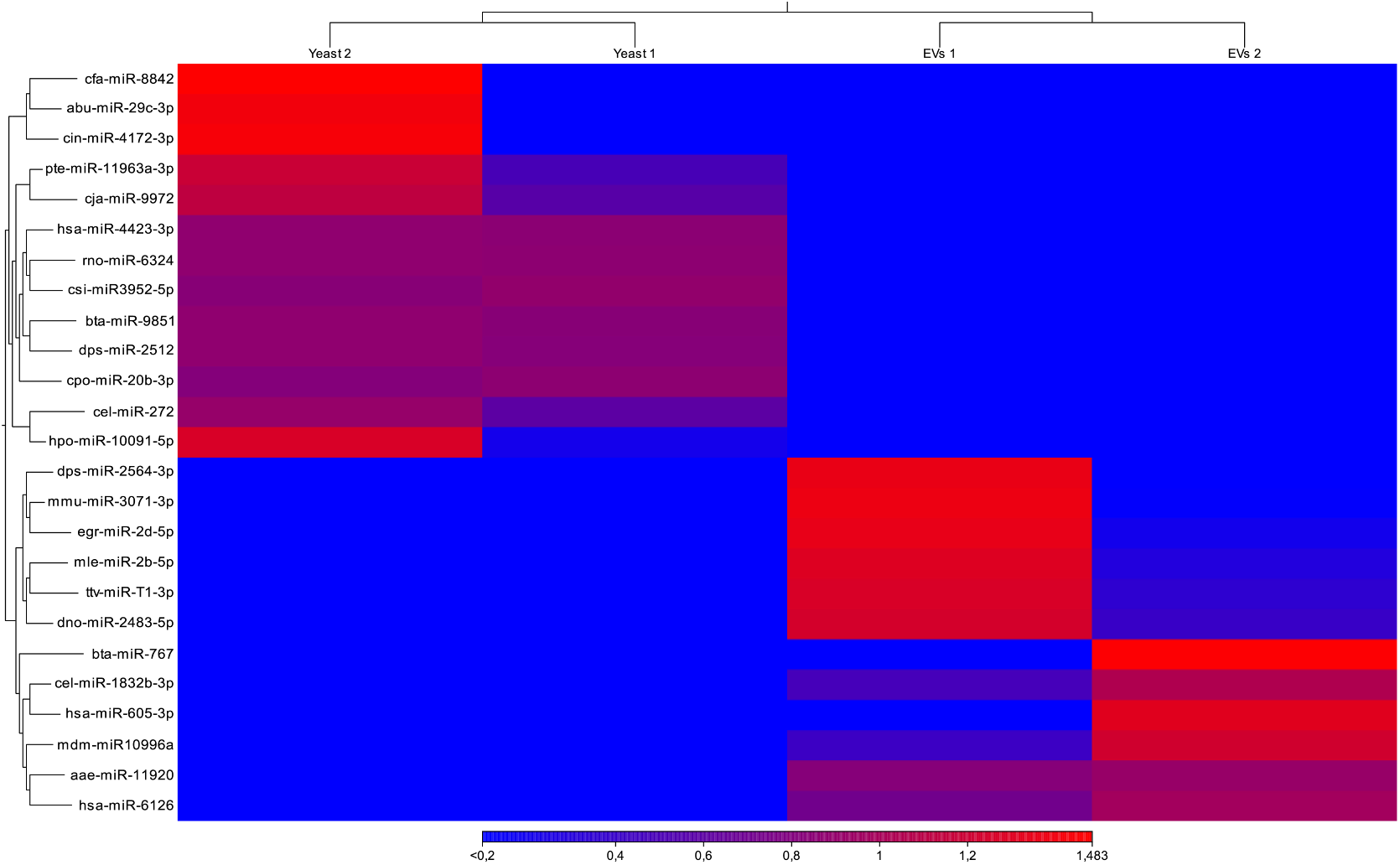
Expression of miRNA-like in yeast and the extracellular vesicles (EVs) of *Candida haemulonii* var *vulnera*. Heatmap illustrating the expression levels of miRNA-like sequences identified in both yeast cells and EVs of *Candida haemulonii* var. *vulnera* 1112/2016

Furthermore, GO analysis revealed significant expression of miRNA-like involved in mitochondrial components, particularly members of the electron transport chain, sugar metabolism, stress response, and virulence. (Fig. 5).

**FIG 5.**
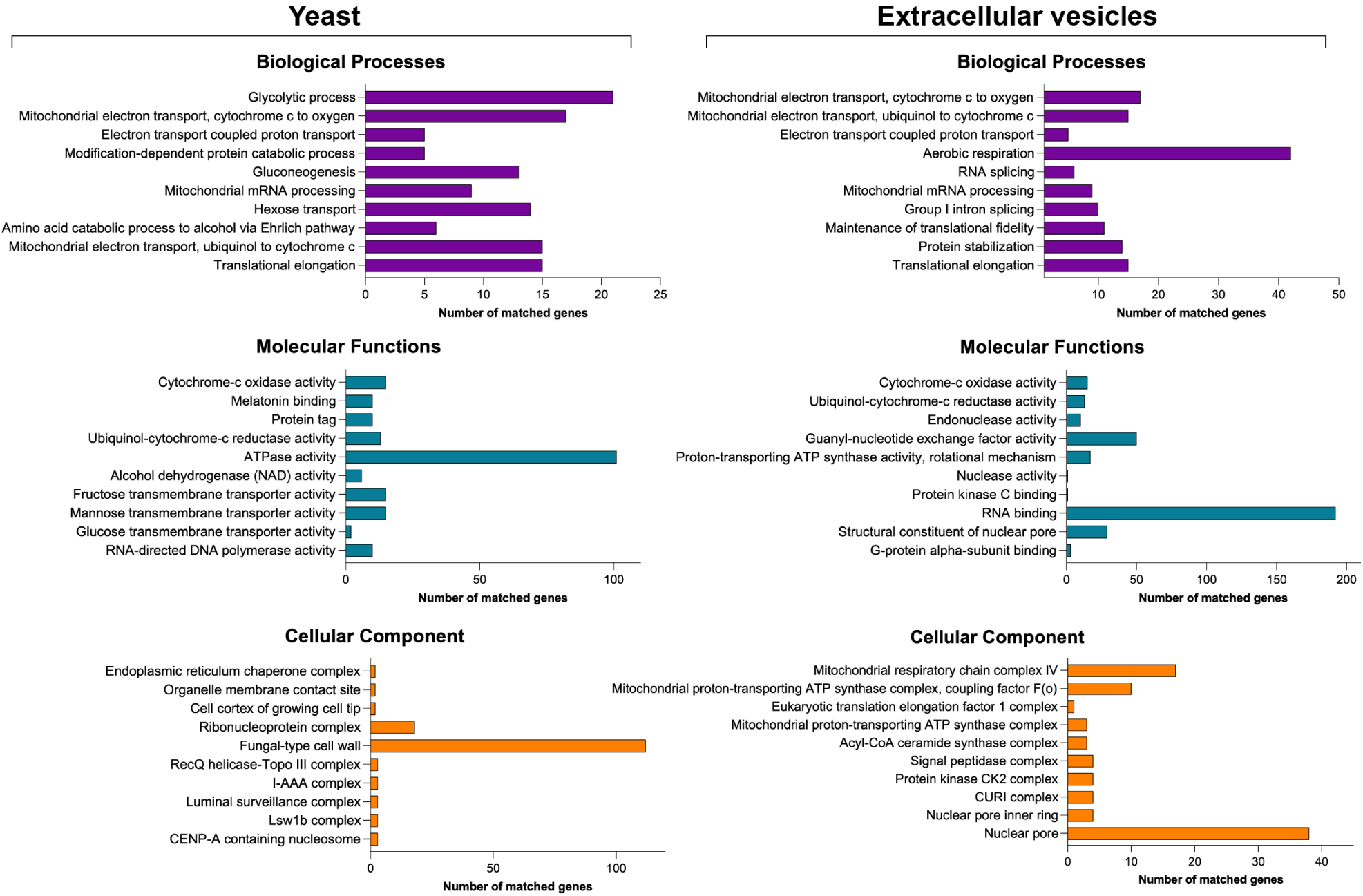
Gene ontology function profile of proteins corresponding to high-coverage RNA sequences identified in miRNA-enriched fractions from extracellular vesicles (EVs) and yeast cells of *Candida haemulonii* var. *vulnera*. The x-axis indicates the number of hits identified for each term in the gene ontology function profile of proteins associated with high-coverage RNA sequences isolated from miRNA-enriched fractions of both extracellular vesicles and yeast cells of *Candida haemulonii* var*. vulnera*.

### Network of significant miRNA interactions from the transcriptomic analysis

Based on the insights gained from protein interactomes, we proceeded to compare the presence of these genes with the genomic data of *C. albicans*. Our analysis of a distinct set of central genes prominently expressed in *Candida haemulonii* var. *vulnera,* particularly those involved in ribosomal biogenesis and function. Genes including RPL5, RPL3, RPL25, RPS3, RPP0, and MRT4 are noteworthy, and they play crucial roles in protein synthesis and cellular metabolism. Additionally, we observed significant expression of chaperone genes such as CCT7, CCT5, and SSZ1, indicating their involvement in protein folding and cellular homeostasis. Conversely, the hub genes identified in EVs exhibited associations with metabolic processes essential for energy production, specifically the citric acid cycle and the electron transport chain. These findings underscore the differential molecular strategies employed by *Candida haemulonii* var. *vulnera* to adapt and thrive in diverse environmental conditions, implicating these pathways in fungal pathogenesis and survival (Fig. 6).

**FIG 6.**
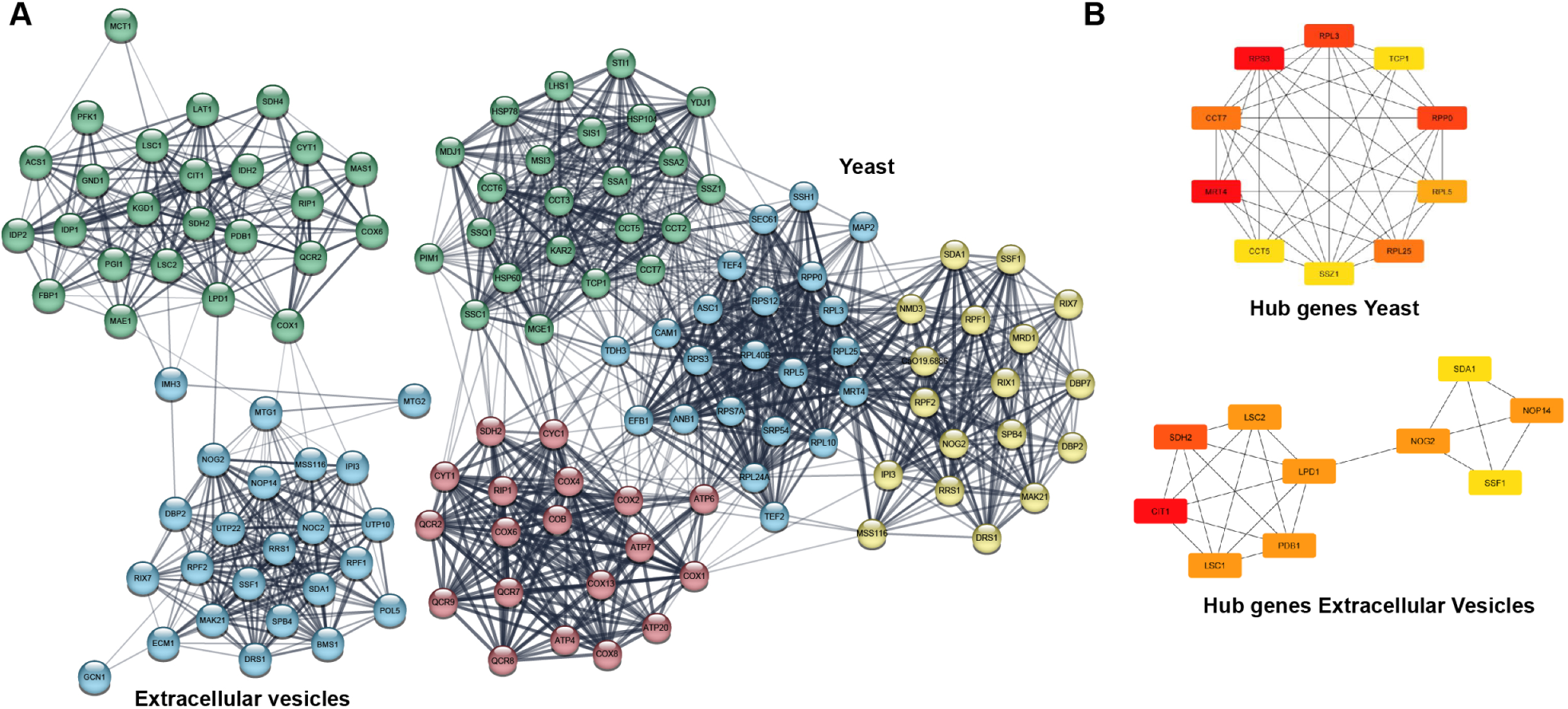
Key genes and miRNA-mediated interactions in *Candida haemulonii* var. *vulnera*. A) Molecular interaction networks of target genes associated with GO functional pathways enriched by miRNA-like from *Candida haemulonii* var. *vulnera*. Networks were constructed using STRING and visualized in Cytoscape based on co-expression analysis, with *Candida albicans* chosen as the reference organism (NCBI taxonomy Id: 237561). **B) Hub genes within the molecular interaction networks observed in *Candida haemulonii var. vulnera* yeast cells and EVs.**

## DISCUSSION

Our investigations into *C. haemulonii* var. *vulnera* revealed a protein profile that shares similarities with previous findings for *C. auris*, which is expected given their phylogenetic and phenotypic closeness [53]. Both yeast cells and EVs from *C. haemulonii* var. *vulnera* exhibited a diverse array of proteins, including ribosomal, ADP/ATP transporters, cell wall components, transcription factors, and various enzymes and subunits. Among the most abundant proteins identified in *C. haemulonii* var. *vulnera* EVs were those involved in starch and sucrose metabolism, protein processing in the endoplasmic reticulum, and antioxidant activities [36]. Similarly, the proteomic analysis for *C. auris* showed parallels with our findings, highlighting various protein classes associated with cell wall integrity, structural components, and specific membrane transporters [54].

The *Candida haemulonii* species complex comprises pathogenic yeasts that have become significant global pathogens [52]. Clinical isolates of *C. haemulonii* have been increasingly reported in recent years due to their capacity to cause invasive infections in immunocompromised individuals [55], inherent resistance to multiple antifungal agents [23], and ability to thrive in diverse environmental conditions [56, 57]. Previously, infections caused by *C. haemulonii* var*. vulnera* were reported in Argentina, Peru, India, and Brazil [2]. The rise and persistence of these cases underscore the critical need for prompt identification and appropriate administration of antifungal therapy that targets these emerging, highly resistant pathogens [58].

Fungal EVs play a crucial role in coordinating fungal communities, enhancing pathogenicity, and influencing interactions with the environment [59]. Previous studies have highlighted EVs’ involvement as immunoregulatory molecules capable of stimulating cytokine production and promoting pathogen infection [60, 61]. For instance, EVs from *Cryptococcus neoformans,* when internalized by macrophages, induce cytokine production and enhance antifungal activity through reactive oxygen species (ROS) production [20, 62]. Understanding the cargo of EVs in these contexts could provide insights into the underlying principles of fungal pathogenicity [63]. Research focusing on *Candida* species has revealed a diverse array of macromolecular classes in EVs [33], with investigations indicating that vesicle content significantly depends on environmental conditions and temporal factors [64].

The selective export of proteins via EVs represents a sophisticated strategy for fungal persistence [65]. It has been noted that the protein composition of *C. albicans* EVs varies depending on the strain morphology [66]. Specifically, EVs derived from hyphal cells contain more virulence-associated proteins, which exhibit cytotoxic effects on human macrophages and induce the release of cytokines such as TNF-alpha [67]. In our study, we identified the presence of the proteasome system as one of the distinctive proteins in EVs from *C. haemulonii* var. *vulnera*. Previous research on *C. albicans* has highlighted the proteasome’s role in regulating protein homeostasis and its essential contribution to virulence [68]. Moreover, the proteasome complex is crucial for managing morphological transitions and maintaining proteostasis under various physiological and stress conditions[69]. These functions are conserved across other *Candida* species, including *C. dubliniensis*, *C. tropicalis*, *C. krusei*, and *C. parapsilosis* [70, 71]. We propose that EV-mediated protein export plays a significant role in enabling *Candida haemulonii* var. *vulnera* yeast to adapt to diverse host environments [72]. The exportation of the proteasome complex likely orchestrates morphological transitions, thereby enhancing virulence and facilitating rapid adaptation in this strain [73]. Additionally, studies have demonstrated the essential role of ubiquitin, an integral component of the proteasome complex, in the growth and virulence of *C. albicans* [74–77].

During pathogen-host interactions, certain proteins play pivotal roles in fungal survival, crucial for establishing infection by evading host defenses, surviving in low-oxygen tissues, undergoing morphological transitions, and releasing virulence factors [33, 66]. One such protein, BMH1, belongs to the 14-3-3 protein family and is involved in adaptive signaling [78]. The direct interactome of 14-3-3 proteins suggests their role in promoting cellular survival under conditions like oxidative stress, DNA damage, or nutrient scarcity [79]. In *Aspergillus flavus*, the absence of BMH1 led to a moderate decrease in vegetative growth and alterations in morphology and secondary metabolic pathways [80]. Our findings regarding *C. haemulonii* var. *vulnera* indicated that BMH1 exhibits functional diversity related to metabolism maintenance, thermal responsiveness, cytoplasmic architecture, and cell wall remodeling. The export of BMH1 via EVs underscores its crucial role in fungal adaptation, supported by existing literature on fungal biology [81].

Another protein exhibiting higher expression in both fractions is TEF1, which plays a crucial role in regulating translation efficiency and coordinating protein synthesis essential for cell viability [82]. In *Leishmania donovani*, TEF1 has been implicated in reducing the activity of infected macrophages [83]. Increased expression of TEF1 in *Pichia pastoris* indicates adaptation to growth under limited carbon source conditions [84]. In *C. albicans*, TEF1 belongs to a group of cytoplasmic proteins frequently exported via EVs, where they serve signaling functions to regulate fungal community dynamics or modulate host responses [85, 86]. TEF-1alpha has been proposed as a genetic marker for *Candida* species identification due to its highly conserved sequence across strains [87]. In *C. haemulonii* var. *vulnera*, TEF1 is integral to numerous metabolic processes, cell division, and responsiveness to the microenvironment [88].

*C. haemulonii* var. *vulnera* relies on acquiring sufficient nutrients to maintain cellular balance, with glucose serving as a crucial substrate for stability [89]. During infection, the pathogen must adapt to varying glucose levels in different human niches [90, 91]. The CDC19 protein plays a pivotal role as an enzyme that converts phosphoenolpyruvate into pyruvate and ATP during glycolysis [92]. In *C. albicans*, this enzyme has been shown to significantly impact pathogenicity; strains lacking CDC19 expression exhibit exhibited reduced virulence due to their inability to transition morphologically into hyphae within phagocytes [93, 94]. Our finding for *C. haemulonii* var. *vulnera* demonstrates that CDC19 is exported via EVs, consistent with observations in other *Candida* species [95]. Moreover, CDC19 is uniquely associated with inducing a host defense response, illustrating its role in modulating the immune response, driving the yeast-to-yeast transition, facilitating cell wall remodeling, and enhancing species virulence.

PDC11 plays a crucial role in redirecting pyruvate towards fermentation and lipid synthesis [96, 97]. Its function is essential for fungal survival, as demonstrated by its deletion in *Saccharomyces cerevisiae* resulting in lethality [98, 99]. The presence of PDC in *C*.

*albicans* underscores its significant metabolic flexibility, allowing for the switch between fermentative and oxidative metabolism [100]. This adaptability enables *Candida* species to thrive in organs and tissues where oxygen is limited, facilitating infiltrative growth [101, 102]. Our findings with *C. haemulonii* var. *vulnera* indicate that the export of PDC through EVs contributes to several adaptive and virulent mechanisms of the cell. This includes the production of the biofilm matrix [103] and the ability to cause both superficial and deep infections [104].

In the context of infection, fungi employ miRNA transport via EVs as a sophisticated survival strategy, utilizing endogenous miRNA-like to modulate gene expression in host cells [42, 43]. These miRNAs act as versatile regulators, capable of either activating or repressing the translation of target messenger RNA molecules within the host [46, 105]. Our study aligns with previous findings in human pathogens such as *C. neoformans*, *Paracoccidiodes brasiliensis*, and *C. albicans*. Drawing from the model organism *S. cerevisiae* [106], our results suggest that EVs from *C. haemulonii* var. *vulnera* transport miRNA-like involved in essential biological processes critical for yeast homeostasis. These processes include functions responsible for mRNA degradation, translation inhibition, or silencing mechanisms.

Our research findings also uncovered the expression of homologous miRNA-like identified in humans and other species, which could play a pivotal role in *C. haemulonii* pathogenesis. For instance, Hsa-miR-4423-3p encodes the *ZNF* gene in humans, a member of the zinc finger protein (ZNF) family known for diverse functions such as DNA recognition, RNA packaging, transcriptional activation, apoptosis regulation, protein folding, assembly, and lipid binding [107]. In *C. neoformans*, ZNF proteins play crucial roles in sexual reproduction, virulence factor production, ionic homeostasis, pathogenesis, and stress resistance [108]. Additionally, the expression of Hpo-miR-10091-5p in nematodes has been shown to manipulate the host immune system, providing insights into the mechanistic framework of RNA transfer [44]. Conversely, the expression of Ttv-miR-T1-3p in viruses allows for modulation of the innate immune response and promotes viral persistence by silencing genes involved in reducing interferon responsiveness and increasing cell proliferation in the presence of this cytokine [109].

The miRNA-like identified in *C. haemulonii* var. *vulnera* yeast cells may contribute to the expression of genes involved in ribosome biogenesis, a process closely linked to growth and proliferation. This mechanism enables yeast to rapidly metabolize sugars through fermentation, sustaining high growth rates and gaining selective advantages in sugar-rich environments [110]. Additionally, it enhances the potential for Polymerase II transcription of other genes, facilitating adaptive responses in the microorganism [111]. In essence, our findings suggest that the expression of ribosomal genes in *C. haemulonii* var. *vulnera* represents an adaptive strategy that allows the organism to maintain a rapid growth rate while efficiently utilizing sugars through fermentation survival and proliferation.

Furthermore, the presence of miRNAs such as TCP1, CCT7, and CCT5, conserved across fungal species, enhances the phenotypic and metabolic capabilities of *C. haemulonii* by facilitating adaptation to environmental stressors [112, 113]. Previous research has shown the CCT complex modulates tolerance and resistance to echinocandins in *C. albicans*, indicating functions beyond maintaining cell wall integrity [114]. These findings suggest that *C. haemulonii* expresses molecules crucial for growth and survival in challenging environments, potentially contributing to its adaptability across diverse host environments [1, 72].Our research revealed that the most abundant transcripts found in *C. haemulonii* var. *vulnera* EVs, with annotated functions, corresponded to proteins involved in the electron transport chain and citric acid cycle, underscoring the fungal dependence on nutrition for competitive advantage. Previous results on *C. albicans* have demonstrated a complex reprogramming of fungal metabolic networks to utilize alternative carbon sources and induce proteins that defend against oxidative and thermal stress [115].

Damage or mutations in the electron transport chain observed in *C. auris* and *C. albicans* have been associated with reduced sensitivity to antifungal agents, including resistance to first-line drugs such as fluconazole [116, 117]. Our findings indicate that *C. haemulonii* var. *vulnera* transports miRNAs via EVs that are associated with mitochondrial components crucial for the virulence of pathogenic fungi. This transport mechanism may be critical during infection, facilitating metabolic crosstalk between the citric acid cycle and oxidative phosphorylation pathways, which are pivotal for the development of pathogenesis [115].

Our findings underscore the critical role of adaptable mechanisms in responding to varying nutrient availability across different host niches, which remains to be fully elucidated for understanding the competence, resistance, and survival strategies of *C. haemulonii* var. *vulnera*. Given the multifaceted roles of EVs in fungal response to stress, virulence, and resistance, it is essential to identify potential therapeutic targets. This may involve targeting specific metabolic pathways, inhibiting critical proteins or miRNAs associated with fungal persistence, thereby exploiting unique genetic vulnerabilities in *C. haemulonii*.

## DATA AVAILABILITY STATEMENT

The raw data supporting the conclusions of this article will be made available by the authors without undue reservation.

## CLINICAL TRIALS

Not applicable.

## ETHICS APPROVAL

Not applicable.

## REFERENCES

1. Cendejas-Bueno, E., et al., Reclassification of the Candida haemulonii Complex as Candida haemulonii (C. haemulonii Group I), C. duobushaemulonii sp. nov. (C. haemulonii Group II), and C. haemulonii var. vulnera var. nov.: Three Multiresistant Human Pathogenic Yeasts. Journal of Clinical Microbiology, 2012. 50(11): p. 3641–3651.

2. Francisco, E.C., A.W. de Jong, and A.L. Colombo, Candida haemulonii Species Complex: A Mini-review. Mycopathologia, 2023. 188(6): p. 909–917.

3. Coles, M., K. Cox, and A. Chao, Candida haemulonii: An emerging opportunistic pathogen in the United States? IDCases, 2020. 21: p. e00900.

4. Ben-Ami, R., et al., Multidrug-Resistant Candida haemulonii and C. auris, Tel Aviv, Israel. Emerging Infectious Diseases, 2017. 23(2).

5. Santos, S.B.D., et al., Presence of Candida spp. and candidiasis in liver transplant patients. An Bras Dermatol, 2018. 93(3): p. 356–361.

6. Gade, L., et al., Understanding the Emergence of Multidrug-Resistant Candida: Using Whole-Genome Sequencing to Describe the Population Structure of Candida haemulonii Species Complex. Frontiers in Genetics, 2020. 11.

7. Rodrigues, L.S., et al., First Genome Sequences of Two Multidrug-Resistant Candida haemulonii var. vulnera Isolates From Pediatric Patients With Candidemia. Frontiers in Microbiology, 2020. 11.

8. Lima, S.L., L. Rossato, and A. Salles de Azevedo Melo, Evaluation of the potential virulence of Candida haemulonii species complex and Candida auris isolates in Caenorhabditis elegans as an in vivo model and correlation to their biofilm production capacity. Microbial Pathogenesis, 2020. 148: p. 104461.

9. Lima, S.L., et al., Increasing Prevalence of Multidrug-Resistant Candida haemulonii Species Complex among All Yeast Cultures Collected by a Reference Laboratory over the Past 11 Years. Journal of Fungi, 2020. 6(3): p. 110.

10. Joffe, L.S., et al., Potential Roles of Fungal Extracellular Vesicles during Infection. mSphere, 2016. 1(4).

11. de Toledo Martins, S., et al., Extracellular Vesicles in Fungi: Composition and Functions. 2018. 422: p. 45–59.

12. Liebana-Jordan, M., et al., Extracellular Vesicles in the Fungi Kingdom. International Journal of Molecular Sciences, 2021. 22(13): p. 7221.

13. Brown, L., et al., Through the wall: extracellular vesicles in Gram-positive bacteria, mycobacteria and fungi. Nature Reviews Microbiology, 2015. 13(10): p. 620–630.

14. Alves, L.R., et al., Extracellular Vesicle-Mediated RNA Release in Histoplasma capsulatum. mSphere, 2019. 4(2).

15. Bitencourt, T.A., et al., Fungal Extracellular Vesicles Are Involved in Intraspecies Intracellular Communication. mBio, 2022.

16. Freitas, M.S., et al., Fungal Extracellular Vesicles as Potential Targets for Immune Interventions. mSphere, 2019. 4(6).

17. Honorato, L., et al., Extracellular Vesicles Regulate Biofilm Formation and Yeast-to-Hypha Differentiation in Candida albicans. mBio, 2022. 13(3).

18. Marina, C.L., et al., Nutritional Conditions Modulate C. neoformans Extracellular Vesicles’ Capacity to Elicit Host Immune Response. Microorganisms, 2020. 8(11): p. 1815.

19. Oliveira, B.T.M., et al., Extracellular Vesicles from Candida haemulonii var. vulnera Modulate Macrophage Oxidative Burst. Journal of Fungi, 2023. 9(5): p. 562.

20. Oliveira, D.b.L., et al., Extracellular Vesicles from Cryptococcus neoformans Modulate Macrophage Functions. Infection and Immunity, 2010. 78(4): p. 1601–1609.

21. Rodrigues, M.L., et al., Extracellular Vesicles Produced by Cryptococcus neoformans Contain Protein Components Associated with Virulence. Eukaryotic Cell, 2008. 7(1): p. 58–67.

22. Voelz, K., et al., ‘Division of labour’ in response to host oxidative burst drives a fatal Cryptococcus gattii outbreak. Nature Communications, 2014. 5(1).

23. Ramos, L.S., et al., Candida haemulonii complex: species identification and antifungal susceptibility profiles of clinical isolates from Brazil. Journal of Antimicrobial Chemotherapy, 2015. 70(1): p. 111–115.

24. Cao, C., et al., Candida haemulonii Species Complex: Emerging Fungal Pathogens of the Metschnikowiaceae Clade. Zoonoses, 2023. 3(1).

25. Allert, S., et al., From environmental adaptation to host survival: Attributes that mediate pathogenicity of Candida auris. Virulence, 2022. 13(1): p. 191–214.

26. Casadevall, A., et al., On the Emergence of Candida auris: Climate Change, Azoles, Swamps, and Birds. mBio, 2019. 10(4).

27. Aguilar-Pontes, M.V., R.P. de Vries, and M. Zhou, (Post-)genomics approaches in fungal research. Brief Funct Genomics, 2014. 13(6): p. 424–39.

28. Hautbergue, T., et al., From genomics to metabolomics, moving toward an integrated strategy for the discovery of fungal secondary metabolites. Nat Prod Rep, 2018. 35(2): p. 147–173.

29. Altelaar, A.F.M., J. Munoz, and A.J.R. Heck, Next-generation proteomics: towards an integrative view of proteome dynamics. Nature Reviews Genetics, 2012. 14(1): p. 35–48.

30. Kim, Y., M.P. Nandakumar, and M.R. Marten, Proteomics of filamentous fungi. Trends in Biotechnology, 2007. 25(9): p. 395–400.

31. Ma, Y., et al., Integrated proteomics and metabolomics analysis of tea leaves fermented by Aspergillus niger, Aspergillus tamarii and Aspergillus fumigatus. Food Chemistry, 2021. 334: p. 127560.

32. Souza, J.A.M., et al., Characterization of Aspergillus fumigatus Extracellular Vesicles and Their Effects on Macrophages and Neutrophils Functions. Frontiers in Microbiology, 2019. 10.

33. Jacobsen, M.D., et al., Specificity of the osmotic stress response in Candida albicans highlighted by quantitative proteomics. Scientific Reports, 2018. 8(1).

34. El-Baz, A.M., et al., Back to Nature: Combating Candida albicans Biofilm, Phospholipase and Hemolysin Using Plant Essential Oils. Antibiotics, 2021. 10(1): p. 81.

35. Kołaczkowska, A. and M. Kołaczkowski, Drug resistance mechanisms and their regulation in Candida non-albicans species. Journal of Antimicrobial Chemotherapy, 2016. 71(6): p. 1438–1450.

36. Zamith-Miranda, D., et al., Comparative Molecular and Immunoregulatory Analysis of Extracellular Vesicles from Candida albicans and Candida auris. mSystems, 2021. 6(4).

37. Vargas, G., et al., Compositional and immunobiological analyses of extracellular vesicles released by Candida albicans. Cellular Microbiology, 2015. 17(3): p. 389–407.

38. de Vries, R.P., et al., Comparative genomics reveals high biological diversity and specific adaptations in the industrially and medically important fungal genus Aspergillus. Genome Biol, 2017. 18(1): p. 28.

39. Wang, Z., M. Gerstein, and M. Snyder, RNA-Seq: a revolutionary tool for transcriptomics. Nature Reviews Genetics, 2009. 10(1): p. 57–63.

40. Bitencourt, T.A., et al., The RNA Content of Fungal Extracellular Vesicles: At the “Cutting-Edge” of Pathophysiology Regulation. Cells, 2022. 11(14): p. 2184.

41. Valadi, H., et al., Exosome-mediated transfer of mRNAs and microRNAs is a novel mechanism of genetic exchange between cells. Nature Cell Biology, 2007. 9(6): p. 654–659.

42. Munhoz da Rocha, I.F., et al., Cross-Kingdom Extracellular Vesicles EV-RNA Communication as a Mechanism for Host–Pathogen Interaction. Frontiers in Cellular and Infection Microbiology, 2020. 10.

43. Peres da Silva, R., et al., Comparison of the RNA Content of Extracellular Vesicles Derived from Paracoccidioides brasiliensis and Paracoccidioides lutzii. Cells, 2019. 8(7): p. 765.

44. Buck, A.H., et al., Exosomes secreted by nematode parasites transfer small RNAs to mammalian cells and modulate innate immunity. Nature Communications, 2014. 5(1).

45. Zhang, T., et al., Cotton plants export microRNAs to inhibit virulence gene expression in a fungal pathogen. Nature Plants, 2016. 2(10).

46. Cai, Q., et al., Plants send small RNAs in extracellular vesicles to fungal pathogen to silence virulence genes. Science, 2018. 360(6393): p. 1126-1129.

47. Vallejo, M.C., et al., The Pathogenic Fungus Paracoccidioides brasiliensis Exports Extracellular Vesicles Containing Highly Immunogenic -Galactosyl Epitopes. Eukaryotic Cell, 2011. 10(3): p. 343–351.

48. Ferreira, G.A., et al., Proteomic analysis of exosomes secreted during the epithelial-mesenchymal transition and potential biomarkers of mesenchymal high-grade serous ovarian carcinoma. Journal of Ovarian Research, 2023. 16(1).

49. Shannon, P., et al., Cytoscape: A Software Environment for Integrated Models of Biomolecular Interaction Networks. Genome Research, 2003. 13(11): p. 2498–2504.

50. Bader, G.D. and C.W.V. Hogue, An automated method for finding molecular complexes in large protein interaction networks. BMC Bioinformatics, 2003. 4(1): p. 2.

51. Chin, C.-H., et al., cytoHubba: identifying hub objects and sub-networks from complex interactome. BMC Systems Biology, 2014. 8(S4).

52. Gómez-Gaviria, M., et al., Candida haemulonii Complex and Candida auris: Biology, Virulence Factors, Immune Response, and Multidrug Resistance. Infection and Drug Resistance, 2023. Volume 16: p. 1455–1470.

53. Desoubeaux, G., et al., Overview about Candida auris: What’s up 12 years after its first description? Journal of Medical Mycology, 2022. 32(2): p. 101248.

54. Zamith-Miranda, D., et al., Multi-omics Signature of Candida auris, an Emerging and Multidrug-Resistant Pathogen. mSystems, 2019. 4(4).

55. Xu, Y.-C., et al., Distribution and Antifungal Susceptibility of Candida Species Causing Candidemia in China: An Update From the CHIF-NET Study. The Journal of Infectious Diseases, 2020. 221(Supplement_2): p. S139-S147.

56. Fonseca, M.S., et al., First description of Candida haemulonii infecting a snake Boa constrictor: Molecular, pathological and antifungal sensitivity characteristics. Microbial Pathogenesis, 2023. 180: p. 106164.

57. Nosanchuk, J., et al., Candida haemulonii complex, an emerging threat from tropical regions? PLOS Neglected Tropical Diseases, 2023. 17(7): p. e0011453.

58. Ramos, L.S., et al., The Threat Called Candida haemulonii Species Complex in Rio de Janeiro State, Brazil: Focus on Antifungal Resistance and Virulence Attributes. Journal of Fungi, 2022. 8(6): p. 574.

59. Nenciarini, S. and D. Cavalieri, Immunomodulatory Potential of Fungal Extracellular Vesicles: Insights for Therapeutic Applications. Biomolecules, 2023. 13(10): p. 1487.

60. Brauer, V.S., et al., Extracellular Vesicles from Aspergillus flavus Induce M1 Polarization In Vitro. mSphere, 2020. 5(3).

61. Almeida, F., et al., Galectin-3 impacts Cryptococcus neoformans infection through direct antifungal effects. Nature Communications, 2017. 8(1).

62. Nielsen, K., et al., Cryptococcus neoformans-Derived Microvesicles Enhance the Pathogenesis of Fungal Brain Infection. PLoS ONE, 2012. 7(11): p. e48570.

63. Reis, F.C.G., et al., Small Molecule Analysis of Extracellular Vesicles Produced by Cryptococcus gattii: Identification of a Tripeptide Controlling Cryptococcal Infection in an Invertebrate Host Model. Frontiers in Immunology, 2021. 12.

64. Zamith-Miranda, D., et al., Transcriptional and translational landscape of Candida auris in response to caspofungin. Computational and Structural Biotechnology Journal, 2021. 19: p. 5264–5277.

65. Albuquerque, P.C., et al., Vesicular transport in Histoplasma capsulatum: an effective mechanism for trans-cell wall transfer of proteins and lipids in ascomycetes. Cellular Microbiology, 2008. 10(8): p. 1695–1710.

66. Gil-Bona, A., et al., Candida albicans cell shaving uncovers new proteins involved in cell wall integrity, yeast to hypha transition, stress response and host–pathogen interaction. Journal of Proteomics, 2015. 127: p. 340–351.

67. Martínez-López, R., et al., Candida albicans Hyphal Extracellular Vesicles Are Different from Yeast Ones, Carrying an Active Proteasome Complex and Showing a Different Role in Host Immune Response. Microbiology Spectrum, 2022. 10(3).

68. Liu, T.-B. and C. Xue, The Ubiquitin-Proteasome System and F-box Proteins in Pathogenic Fungi. Mycobiology, 2018. 39(4): p. 243–248.

69. Cao, C. and C. Xue, More Than Just Cleaning: Ubiquitin-Mediated Proteolysis in Fungal Pathogenesis. Frontiers in Cellular and Infection Microbiology, 2021. 11.

70. Enenkel, C., et al., Intracellular localization of the proteasome in response to stress conditions. Journal of Biological Chemistry, 2022. 298(7): p. 102083.

71. Leach, M.D., et al., Molecular and proteomic analyses highlight the importance of ubiquitination for the stress resistance, metabolic adaptation, morphogenetic regulation and virulence of Candida albicans. Molecular Microbiology, 2011. 79(6): p. 1574–1593.

72. Garcia-Bustos, V., et al., Climate change, animals, and Candida auris: insights into the ecological niche of a new species from a One Health approach. Clinical Microbiology and Infection, 2023. 29(7): p. 858–862.

73. Angiolella, L., et al., Identification of Virulence Factors in Isolates of Candida haemulonii, Candida albicans and Clavispora lusitaniae with Low Susceptibility and Resistance to Fluconazole and Amphotericin B. Microorganisms, 2024. 12(1): p. 212.

74. Liu, W., et al., The Ubiquitin Conjugating Enzyme: An Important Ubiquitin Transfer Platform in Ubiquitin-Proteasome System. International Journal of Molecular Sciences, 2020. 21(8): p. 2894.

75. Yang, D., et al., Candida albicans Ubiquitin and Heat Shock Factor-Type Transcriptional Factors Are Involved in 2-Dodecenoic Acid-Mediated Inhibition of Hyphal Growth. Microorganisms, 2020. 8(1): p. 75.

76. Hossain, S., et al., The Proteasome Governs Fungal Morphogenesis via Functional Connections with Hsp90 and cAMP-Protein Kinase A Signaling. mBio, 2020. 11(2).

77. Mitchell, A.P., et al., The proteasome regulator Rpn4 controls antifungal drug tolerance by coupling protein homeostasis with metabolic responses to drug stress. PLOS Pathogens, 2023. 19(4): p. e1011338.

78. Pennington, K.L., et al., The dynamic and stress-adaptive signaling hub of 14-3-3: emerging mechanisms of regulation and context-dependent protein–protein interactions. Oncogene, 2018. 37(42): p. 5587–5604.

79. Lehtinen, M.K., et al., A Conserved MST-FOXO Signaling Pathway Mediates Oxidative-Stress Responses and Extends Life Span. Cell, 2006. 125(5): p. 987–1001.

80. Ibarra, B.A., et al., The 14-3-3 Protein Homolog ArtA Regulates Development and Secondary Metabolism in the Opportunistic Plant Pathogen Aspergillus flavus. Applied and Environmental Microbiology, 2018. 84(5).

81. Shi, L., et al., 14-3-3 Proteins: a window for a deeper understanding of fungal metabolism and development. World Journal of Microbiology and Biotechnology, 2019. 35(2).

82. Satala, D., et al., Moonlighting Proteins at the Candidal Cell Surface. Microorganisms, 2020. 8(7): p. 1046.

83. Nandan, D., et al., Leishmania EF-1 Activates the Src Homology 2 Domain Containing Tyrosine Phosphatase SHP-1 Leading to Macrophage Deactivation. Journal of Biological Chemistry, 2002. 277(51): p. 50190–50197.

84. Ahn, J., et al., Translation elongation factor 1- gene from Pichia pastoris: molecular cloning, sequence, and use of its promoter. Applied Microbiology and Biotechnology, 2007. 74(3): p. 601–608.

85. Hernáez, M.L., et al., Identification of Candida albicans exposed surface proteins in vivo by a rapid proteomic approach. Journal of Proteomics, 2010. 73(7): p. 1404–1409.

86. Pitarch, A., et al., Sequential Fractionation and Two-dimensional Gel Analysis Unravels the Complexity of the Dimorphic Fungus Candida albicans Cell Wall Proteome. Molecular & Cellular Proteomics, 2002. 1(12): p. 967–982.

87. Pakshir, K., et al., Translation elongation factor 1-alpha gene as a marker for diagnosing of Candida onychomycosis. Current Medical Mycology, 2020.

88. Seweryn, K., et al., Kinetic and thermodynamic characterization of the interactions between the components of human plasma kinin-forming system and isolated and purified cell wall proteins of Candida albicans. Acta Biochimica Polonica, 2015. 62(4): p. 825–835.

89. Rutherford, J.C., et al., Nutrient and Stress Sensing in Pathogenic Yeasts. Frontiers in Microbiology, 2019. 10.

90. Van Ende, M., S. Wijnants, and P. Van Dijck, Sugar Sensing and Signaling in Candida albicans and Candida glabrata. Frontiers in Microbiology, 2019. 10.

91. Chew, S.Y., W.J.Y. Chee, and L.T.L. Than, The glyoxylate cycle and alternative carbon metabolism as metabolic adaptation strategies of Candida glabrata: perspectives from Candida albicans and Saccharomyces cerevisiae. Journal of Biomedical Science, 2019. 26(1).

92. Wijnants, S., et al., Sugar Phosphorylation Controls Carbon Source Utilization and Virulence of Candida albicans. Frontiers in Microbiology, 2020. 11.

93. Barelle, C.J., et al., Niche-specific regulation of central metabolic pathways in a fungal pathogen. Cellular Microbiology, 2006. 8(6): p. 961–971.

94. Tucey, T.M., et al., Glucose Homeostasis Is Important for Immune Cell Viability during Candida Challenge and Host Survival of Systemic Fungal Infection. Cell Metabolism, 2018. 27(5): p. 988–1006.e7.

95. Karkowska-Kuleta, J., et al., Characteristics of Extracellular Vesicles Released by the Pathogenic Yeast-Like Fungi Candida glabrata, Candida parapsilosis and Candida tropicalis. Cells, 2020. 9(7): p. 1722.

96. Khemacheewakul, J., et al., Validation of mathematical model with phosphate activation effect by batch (R)-phenylacetylcarbinol biotransformation process utilizing Candida tropicalis pyruvate decarboxylase in phosphate buffer. Scientific Reports, 2021. 11(1).

97. Mushtaq, Z. and H. Mukhtar, Process optimization for biosynthesis of pyruvate decarboxylase (PDC) and Neuberg’s ketol (PAC) from a novel Pichia cecembensis through response surface methodology. Annals of Microbiology, 2022. 72(1).

98. Misra, H.S., et al., Pyruvate decarboxylase and thiamine biosynthetic genes are regulated differently by Pdc2 in S. cerevisiae and C. glabrata. Plos One, 2023. 18(6): p. e0286744.

99. Flikweert, M.T., et al., Pyruvate decarboxylase: an indispensable enzyme for growth of Saccharomyces cerevisiae on glucose. Yeast, 1996. 12(3): p. 247–257.

100. Tylicki, A., et al., Comparative study of the activity and kinetic properties of malate dehydrogenase and pyruvate decarboxylase from Candida albicans, Malassezia pachydermatis, and Saccharomyces cerevisiae. Canadian Journal of Microbiology, 2008. 54(9): p. 734–741.

101. Vashist, A., et al., Changes in the mRNA expression of glycolysis-related enzymes of Candida albicans during inhibition of intramitochondrial catabolism under anaerobic condition. Plos One, 2023. 18(4): p. e0284353.

102. Luo, Z., et al., Enhanced pyruvate production in Candida glabrata by carrier engineering. Biotechnology and Bioengineering, 2017. 115(2): p. 473–482.

103. Ramos, L.S., et al., Biofilm Formed by Candida haemulonii Species Complex: Structural Analysis and Extracellular Matrix Composition. Journal of Fungi, 2020. 6(2): p. 46.

104. Chen, X.-F., et al., First two fungemia cases caused by Candida haemulonii var. vulnera in China with emerged antifungal resistance. Frontiers in Microbiology, 2022. 13.

105. Kwon, S., et al., Inside-out: from endosomes to extracellular vesicles in fungal RNA transport. Fungal Biology Reviews, 2020. 34(2): p. 89–99.

106. Peres da Silva, R., et al., Extracellular vesicle-mediated export of fungal RNA. Scientific Reports, 2015. 5(1).

107. Caza, M., et al., The Zinc Finger Protein Mig1 Regulates Mitochondrial Function and Azole Drug Susceptibility in the Pathogenic Fungus Cryptococcus neoformans. mSphere, 2016. 1(1).

108. Li, Y.-H. and T.-B. Liu, Zinc Finger Proteins in the Human Fungal Pathogen Cryptococcus neoformans. International Journal of Molecular Sciences, 2020. 21(4): p. 1361.

109. Teng, Y., et al., Systematic Genome-wide Screening and Prediction of microRNAs in EBOV During the 2014 Ebolavirus Outbreak. Scientific Reports, 2015. 5(1).

110. Kaur, R., I. Castano, and B.P. Cormack, Functional Genomic Analysis of Fluconazole Susceptibility in the Pathogenic Yeast Candida glabrata: Roles of Calcium Signaling and Mitochondria. Antimicrobial Agents and Chemotherapy, 2004. 48(5): p. 1600–1613.

111. Yang, C., et al., mRNA Turnover Protein 4 Is Vital for Fungal Pathogenicity and Response to Oxidative Stress in Sclerotinia sclerotiorum. Pathogens, 2023. 12(2): p. 281.

112. Anand, R., et al., Reprogramming in Candida albicans Gene Expression Network under Butanol Stress Abrogates Hyphal Development. International Journal of Molecular Sciences, 2023. 24(24): p. 17227.

113. Sagini, J.P.N. and R. Ligabue-Braun, Fungal heat shock proteins: molecular phylogenetic insights into the host takeover. The Science of Nature, 2024. 111(2).

114. Caplan, T., et al., Functional Genomic Screening Reveals Core Modulators of Echinocandin Stress Responses in Candida albicans. Cell Reports, 2018. 23(8): p. 2292–2298.

115. Butler, G., et al., Integration of the tricarboxylic acid (TCA) cycle with cAMP signaling and Sfl2 pathways in the regulation of CO2 sensing and hyphal development in Candida albicans. PLOS Genetics, 2017. 13(8): p. e1006949.

116. Zhou, M., et al., Divergent mitochondrial responses and metabolic signal pathways secure the azole resistance in Crabtree-positive and negative Candida species. Microbiology Spectrum, 2024. 12(4).

117. Calderone, R., D. Li, and A. Traven, System-level impact of mitochondria on fungal virulence: to metabolism and beyond. FEMS Yeast Research, 2015. 15(4).

